# Interaction between distinct actin pools controls activity-dependent actin dynamics in the dendritic spine

**DOI:** 10.1101/077933

**Authors:** Oleg O. Glebov, Juan Burrone

## Abstract

Actin cytoskeleton is composed of functionally distinct pools of filamentous (F)-actin defined by their regulatory machinery and dynamics. Although these networks may compete for actin monomers and regulatory factors^1–4^, the interaction between them remains poorly understood. Here, we show that disruption of the labile F-actin pool in neurons by limited actin depolymerization^5,6^ unexpectedly triggers rapid enhancement of the F-actin content at the dendritic spine. Long-term blockade of NMDA-type receptors decreases spine actin polymerization, which is specifically restored by the labile pool ablation. Increase in the spine actin is triggered by blockade of formin-induced actin polymerization in a manner dependent on Arp2/3 complex activity. Finally, limited actin depolymerization increases F-actin levels in a cultured cell line, suggesting the generality of the two-tiered actin dynamics. Based on these findings, we propose a model whereby the labile pool of F-actin controlled by formin restricts the polymerization state of the Arp2/3-regulated stable spine actin, suggesting a feedback principle at the core of cytoskeletal organization in neurons.

**Highlights:**

1. Disruption of labile F-actin by limited depolymerization rapidly increases the synaptic F-actin content;
2. The depolymerization-induced F-actin boost reverses decrease in synaptic F-actin induced by long-term NMDA receptor blockade;
3. Blockade of formin-dependent actin polymerization boosts synaptic F-actin in an Arp2/3-dependent manner;
4. Limited actin depolymerization enhances overall F-actin content in a mammalian cell line.

---

Dynamic polymerization of globular (G) actin into the filamentous (F) form is essential for control of neuronal function, and aberrant actin dynamics have been implicated in neuronal pathology^7–9^. The majority of F-actin in mature neurons resides in a comparatively stable^10^ pool localized at the dendritic spine both in the vicinity and at the synapse, where it regulates postsynaptic structure and function^8,11^ through a treadmilling process dependent on the Arp2/3 branching complex activity^12–15^. Additionally, a labile pool of postsynaptic F-actin has been shown to control postsynaptic receptor trafficking^6,16^, but the molecular identity of this pool has not been established. Dendritic spines therefore possess at least two actin pools with distinct functions, but the putative connexion between them remains unknown.

To test the hypothesis that the labile and the stable pools of F-actin in the spine are functionally linked, we selectively disrupted the former, taking advantage of its sensitivity to a low concentration of the commonly employed actin depolymerizing drug Latrunculin A (LatA)^6,16^. To this end, we treated neurons with various concentrations of LatA overnight and quantified the content of F-actin in the dendritic spine using phalloidin staining and immunocytochemistry for a canonical spine marker Homer. As reported before^10,17^, a high (5uM) concentration of LatA led to a significant drop in the spine F-actin content, consistent with local depolymerization of F-actin. Strikingly, a 100-fold lower (50nM) concentration of LatA did not decrease the spine F-actin content, but rather increased it (Fig. 1a,b).

**Figure 1.**
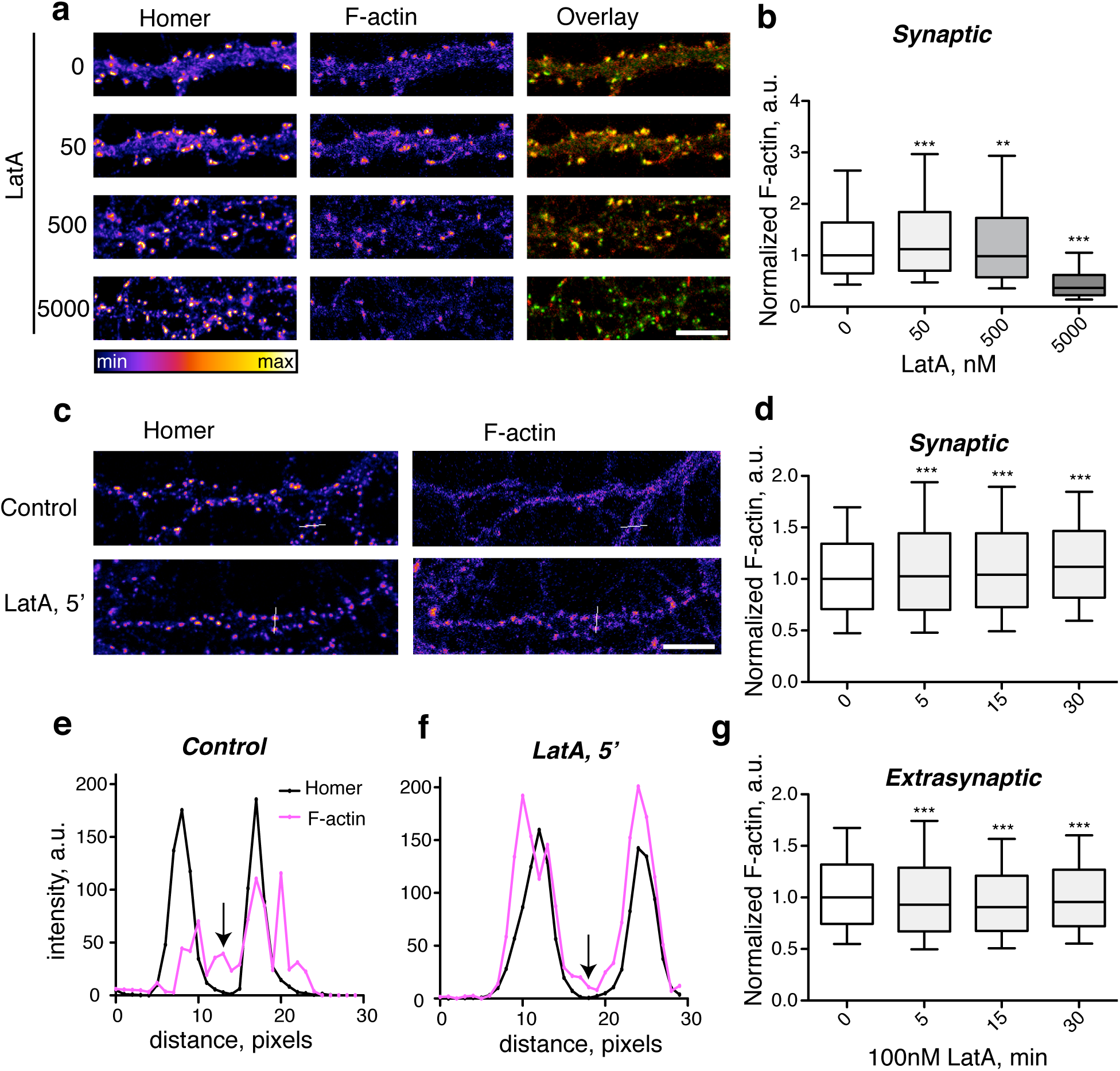
Limited depolymerization of actin results in rapid enhancement of the spine actin content. (**a**) Neurons were incubated with various concentrations of LatA overnight and stained for Homer and Phalloidin. (**b**), quantification of F-actin content in Homer-positive puncta. N=3. (**c**) Neurons were incubated with 100nM LatA and processed as in (a). White lines denote cross-sections of the dendrite, in both cases encompassing two Homer-positive spines separated by a Homer-positive shaft region. (**d**) Quantification of the Phalloidin content in Homer-positive puncta after various times in 100nM LatA. (**e-f**) Intensity profiles of the while lines from the images in (c). (**e**) Untreated sample. Arrow denotes presence of Phalloidin staining in the shaft. (**f**) Sample treated with LatA for 5min. Note the enhanced accumulation of F-actin in the spines and decreased accumulation in the shaft. (**g**) Quantification of the F-actin content for the regions comprising the 0.5um vicinity around Homer-positive puncta. Scale bar, 10um.

Overnight incubation with all of the LatA concentrations resulted in an increase in the levels of Homer, consistent with the strengthening of the synapse following long-term activity blockade^8,18^ (Supplementary Fig. 1a). Given that actin depolymerization may affect neuronal activity through decrease in presynaptic release and activation of postsynaptic receptors^17,19,20^, although F-actin was still comparatively enriched in the synapse relative to the Homer content at 50nM LatA (Supplementary Fig. 1b), we sought to determine whether the increase in F-actin is activity-independent, i.e. whether enlargement of the spine F-actin pool could be uncoupled from the enlargement of the synapse itself. To this end, we tracked the short-term dynamics of neuronal F-actin over 30min following limited depolymerization, using colocalization with Homer as a means for discriminating between spine and non-spine F-actin. The increase in F-actin content colocalizing with the Homer-positive puncta was evident after as little as 5min of incubation with 100nM LatA (Fig. 1c–f), while the content of Homer itself remained unchanged within this time frame (not shown). The increase in the spine F-actin content was mirrored by a decrease in non-synaptic F-actin in the vicinity of the dendritic spine, likely reflecting actin depolymerization in the dendritic shaft and axons (Fig. 1e,g). Furthermore, there was a pronounced loss of F-actin from the cell body, suggesting that depolymerization of F-actin was specifically present in the somatodendritic compartment of the neuron (Supplementary Fig. 1c,d). Enlargement of the spine F-actin pool by limited depolymerization is therefore rapid, independent from neuronal activity and associated with depolymerization of non-synaptic somatodendritic F-actin.

The effect of LatA on actin polymerization is realized by shifting the equilibrium towards depolymerization through sequestering of the G-actin monomers^21^. An alternative actin-depolymerizing drug Cytochalasin D (CytoD) acts in a different manner, preventing the growth of the actin filament by capping its growing end^22^, and does not depolymerize spine F-actin^10^. Upon application of CytoD (1uM), F-actin content in the spine was also rapidly increased (Supplementary Fig. 1e,f). Thus, limited depolymerization of F-actin by two unrelated pharmacological agents acting through two different mechanisms results in an increase in of spine F-actin.

Postsynaptic actin dynamics are acutely controlled by neuronal activity, and this regulation is believed to play a key role in the rapid forms of synaptic plasticity^7^. The role of actin in slower homeostatic forms of synaptic plasticity, on the other hand, remains largely unexplored. To visualize the actin dynamics in the context of the long-term plasticity, we quantified the levels of spine F-actin following a chronic blockade of neuronal activity (48 hours). Blockade of action potential generation by Tetrodotoxin (TTX) resulted in a significant reduction in the amount of F-actin both in the synapse and in the cell body (Fig. 2a,b, Supplementary Fig. 2a,b). Specific blockade of AMPA- and kainate-type glutamate receptors by NBQX had no effect on the levels of F-actin, while blockade of NMDA-type glutamate receptor (NMDAR) by APV had the same effect as the TTX, indicating that signalling through NMDARs was specifically required for spine actin polymerization. 30min treatment with LatA rescued the TTX-induced loss of F-actin from the spines (Fig. 2c,d) but not from the cell body (Supplementary Fig. 2c,d), indicating that depolymerization of the labile pool specifically controlled the activity-dependent actin dynamics at the spine. Importantly, application of TTX significantly increased the overall dendritic levels of F-actin, strongly suggesting that the effect of LatA was not due to redistribution of already polymerized actin, but rather reflected a *bona fide* shift in the polymerization state on neuronal actin (Fig. 2c, Supplementary Fig. 2e).

**Figure 2.**
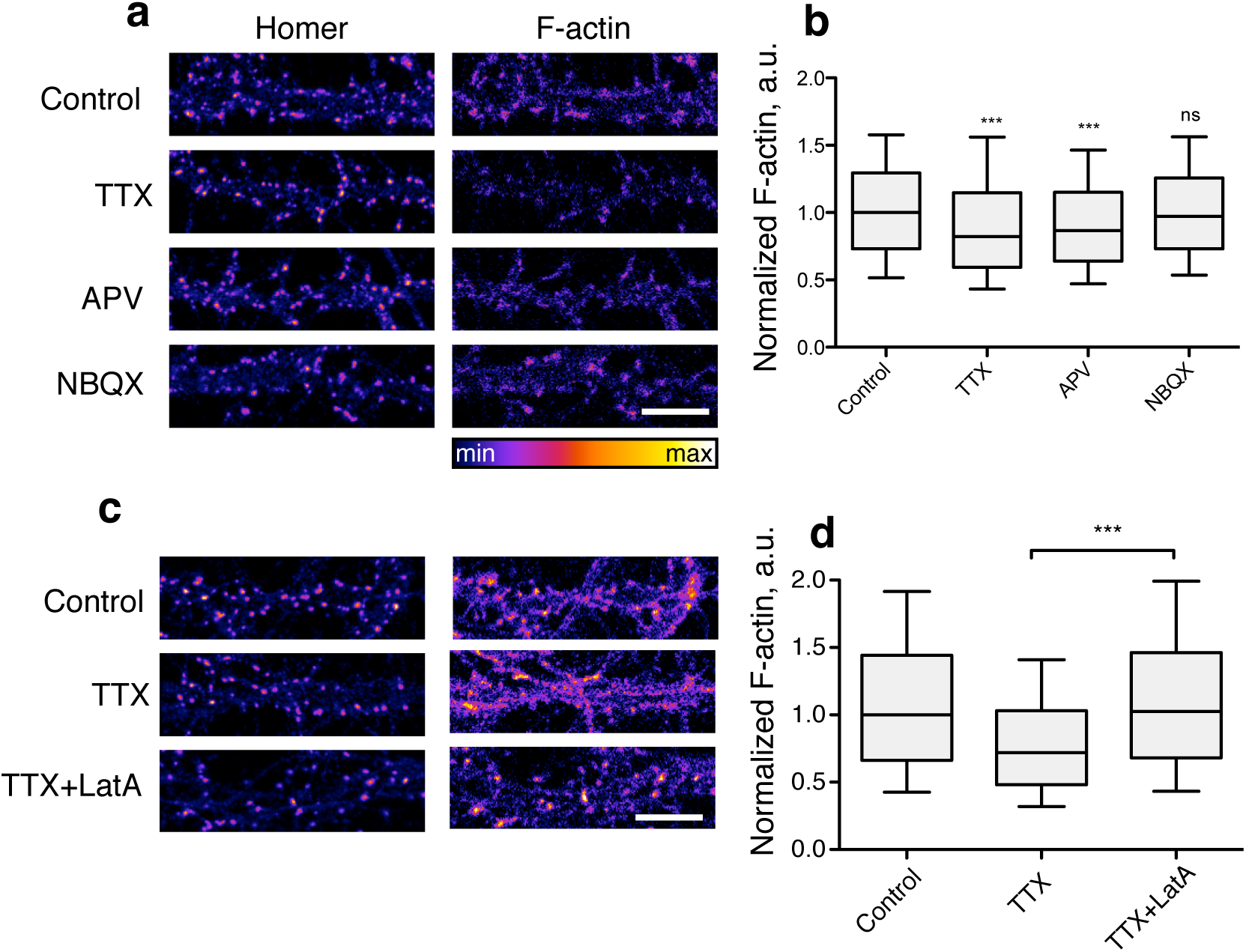
Limited actin depolymerization reverses loss of F-actin from the spine induced by the blockade of NMDA receptors. (**a**) Neurons were incubated for 48h in presence of indicated blockers and stained for F-actin and Homer. (**b**) Quantification of F-actin content in Homer-positive puncta after treatment with indicated blockers. N=3. (**c**) Neurons were incubated for 48h in presence of TTX followed by 30min in presence of LatA, and stained for F-actin and Homer. (**d**) Quantification of f-actin content in Homer-positive puncta after indicated treatment. N=4. Scale bar, 10um.

Our data so far suggested that selective disruption of the labile pool of actin in neurons led to the enhancement of the stable pool. What are the molecular mechanisms defining the distinct identity of these pools? Two competing actin networks in yeast and mammalian cells have been characterized by their dependence on branching complex Arp2/3 and members of a formin family respectively^1–3,23^. Both formins and Arp2/3 are enriched in the spine^24^, where the activity of the latter constitutes a well-established regulatory mechanism controlling local actin dynamics and synaptic function^9,12–14,24^. Formin-dependent but Arp2/3-independent actin network can also be found in the axons where it is sensitive to low concentrations of LatA^5^, hinting that the labile pool observed on our experiments may involve formin activity. Alternatively, an actin-bundling protein Myosin II has been recently shown to antagonize Arp2/3-dependent actin dynamics at the leading edge of migrating cells^4^.

To investigate the putative relationship between these networks in the context of the postsynaptic actin dynamics, we employed pharmacological perturbation of Myosin II, Arp2/3 and formin functions. Blockade of myosin II function by blebbistatin^25^ had no effect on the levels of spine F-actin after 30min, suggesting that release of actin from the myosin-dependent bundled pool did not shift the polymerization/depolymerization balance (not shown). Inhibition of Arp2/3 function by CK-666^26^, on the other hand, decreased the levels of synaptic F-actin and abolished the effect of low LatA, directly confirming requirement for Arp2/3 in maintenance of the spine actin content^14,15^ (Fig. 3e,f). In contrast, blockade of the actin-binding formin homology domain 2 activity by SMIFH2^27^ resulted in an increase in synaptic F-actin, mimicking the effect of low LatA concentration and indicating that formin-dependent actin elongation of F-actin was limiting the spine F-actin levels. Furthermore, the SMIFH2-induced increase in actin levels was abolished by co-application of CK-666, suggesting that formin blockade resulted in and enhancement of the Arp2/3-dependent branching (Fig. 3g,h). Interestingly, inhibition of both formin and Arp2/3 showed no additive effect over inhibition of Arp2/3 alone, suggesting that blockade of Arp2/3-dependent branching also abolished formin-dependent elongation in the spine.

**Figure 3.**
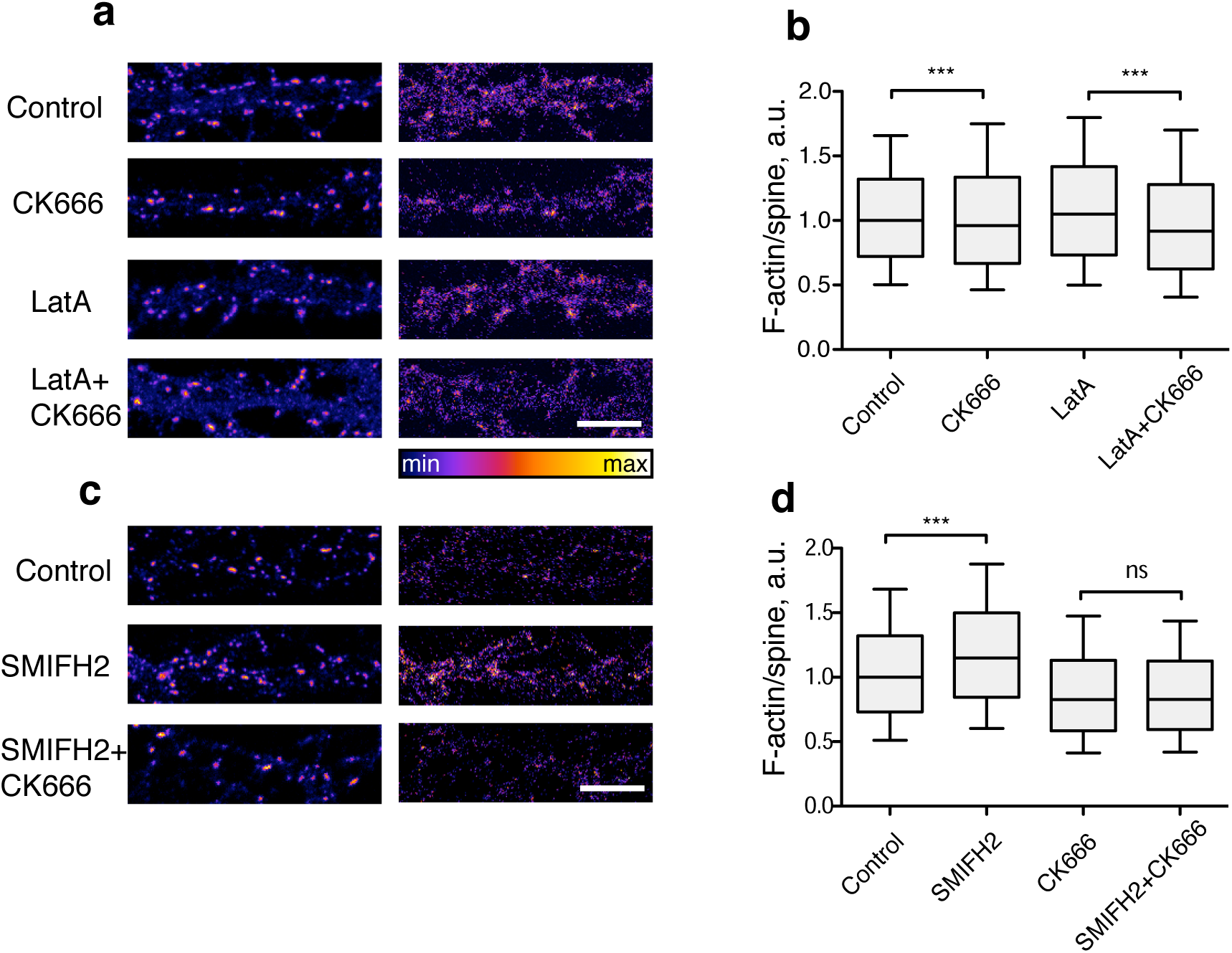
Formin-dependent restriction of the Arp2/3-dependent actin polymerization at the spine. (**a**) Neurons were treated with LatA and/or CK666 for 30min and stained for F-actin and Homer. (**b**) Quantification of F-actin content in Homer-positive puncta after treatment with indicated blockers. N=4. (**c**) Neurons were treated with SMIFH2 and CK666 for 60min and stained for F-actin and Homer. (**d**) Quantification of F-actin content in Homer-positive puncta after treatment with indicated blockers. N=4. Scale bar, 10um.

Finally, to test whether the functional relationship between different pools of F-actin is restricted to neurons or whether it constitutes a general feature of mammalian cells, we probed for potential interplay between pools of actin in another cell type, using the U2OS osteosarcoma cell line^23^. Application of 50nM concentration of LatA for 30min resulted in a decrease of the F/G actin ratio, indicative of a more dynamic actin turnover compared to dendritic spines (Fig. 4a,b). Treatment with an even lower (17nM) concentration of LatA, however, led to an overall increase in the F-actin content comparable to the effect observed in neurons (Fig. 4b). Thus, the labile pool of F-actin may restrict actin dynamics in cells other than neurons.

**Figure 4.**
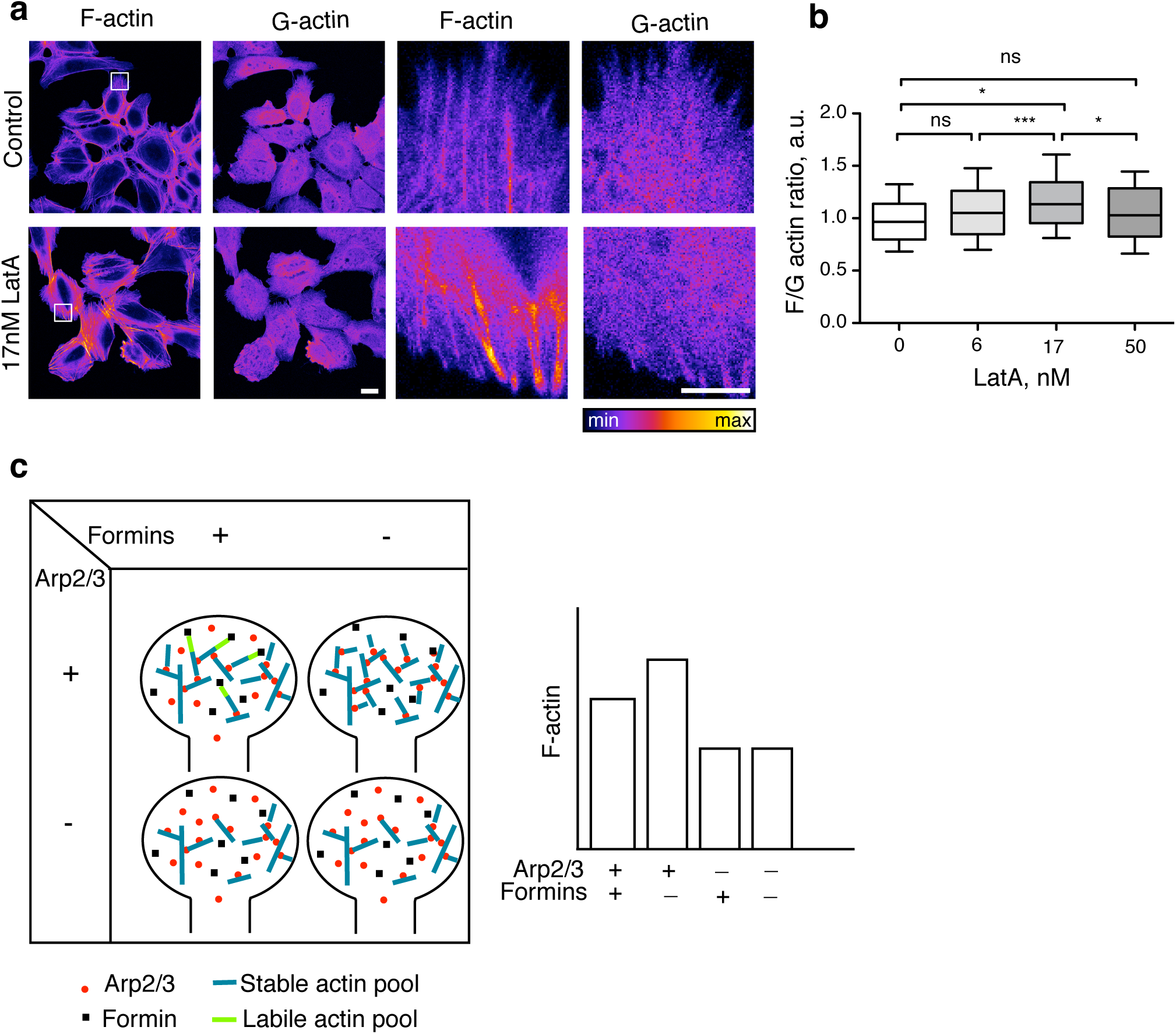
Low concentration of LatA induces actin polymerization in a cell line. (**a**) U2OS cells were incubated with 17nM LatA for 30’ and stained for F-actin and G-actin. (**b**) F/G actin ratio was quantified for 30’ incubations at different LatA concentrations. N=5. (**c**) Proposed model of F-actin organization in the spine. The majority of neuronal F-actin is maintained in the stable spine pool by the Arp2/3 branching activity, along with the minor labile pool elongated by formin. Inhibition of formin triggers an increase in the Arp2/3-dependent actin pool through an unknown mechanism, while blockade of Arp2/3 results in decrease in the Arp2/3 pool. The lack of effect of dual formin-Arp2/3 inhibition compared to Arp2/3 inhibition alone is consistent with formins having no direct role in the spine F-actin polymerization. Scale bar, 25um.

In this study, we have focused on elucidating the relationship between two functionally and dynamically distinct pools of F-actin in neurons. Our data shows that disruption of the labile pool of neuronal F-actin leads to rapid functional changes in the stable pool at the dendritic spine, demonstrating for the first time functional interaction between actin pools in a physiologically relevant primary cell type. Crucially, while previous evidence has focused on a “tug-of-war”-like competition between the actin networks^1–4^, our data indicates that there exists a degree of inequality between them, at least in the context of the dendritic spine.

Our experiments show that the formin-dependent actin elongation makes no major contribution to the polymerization of the spine actin; instead, it appears to restrict the Arp2/3-dependent actin branching activity, as disruption of the labile pool either by low LatA concentration or blockade of formin activity leads to an Arp2/3-dependent accumulation of the F-actin in the spine. Conversely, inhibition of the Arp2/3 function results in a pronounced decrease in spine F-actin, directly confirming the essential role of Arp2/3 in maintaining postsynaptic actin dynamics. Lack of additive effect for Arp2/3 and formin inhibition is consistent with the role for formin in elongation of actin filaments in the spine that have been already initiated by Arp2/3^24^; alternatively, the extra effect of formin inhibition may go unnoticed due to the small size of the formin-dependent pool, as is the case in axons^5^.

This interplay between formin- and Arp2/3-dependent polymerization provides a feedback loop allowing for homeostatic control over the spine actin dynamics, which is likely to be of physiological relevance in the context of long-term alterations of neuronal activity. The marked increase in the spine F-actin content upon inhibition of formin suggests that the regulatory mechanism may be more complex than simple competition for free G-actin^1^, potentially involving other actin-binding proteins such as profilin^2,3^.

It is possible that formin-dependent actin dynamics beyond the spine may also be involved in restriction of Arp2/3 function. This notion is supported by a broad expression profile for formin family members as well as by documented presence of formin-dependent processes elsewhere the neuronal cell^5,24,28,29^. Furthermore, our data (Fig. 4a,b) as well as previously published evidence^1,3^ suggests that the functional interplay between actin pools can be observed beyond the highly specialized environment of the neuronal cell and may therefore represent a general feature of eukaryotic cells. Elucidation of the mechanism allowing for interplay between labile and stable pools of actin will thus warrant further investigation.

## Materials and Methods

### Cell culture

Dissociated hippocampal neuronal cultures were prepared from E18 rat embryos, plated onto poly-L-lysine-coated glass coverslips and maintained according to the standard Banker method. All experiments involving neurons were carried out at 21–27 days in vitro. U2OS cells were grown according to a standard culture protocol. Cells were plated onto 13mm round glass coverslips (thickness 1.0) placed in 35mm Petri dishes (4/dish).

### Reagents

Cell culture media was from Invitrogen. Poly-L-lysine was from Sigma. Primary antibodies raised against the following antigens were used: Homer (160003, Synaptic Systems). Alexa Fluor 488-conjugated anti-rabbit secondary antibody was from Jackson Immunoresearch (USA). Alexa Fluor 568 or 647-conjugated Phalloidin were from Molecular Probes. APV, NBQX, TTX, LIMKi3 were from Tocris (UK). Latrunculin A, CK-666, Cytochalasin D, SMIFH2 and Blebbistatin were from Sigma-Aldrich (UK). Alexa Fluor 594-conjugated DNAse I was from Thermo Fisher.

### Fixed cell imaging

After treatment, coverslips were fixed with 4%PFA in PBS for 15–20min at room temperature (RT) and permeabilized in 0.2%Triton-X100 in PBS supplemented with 5% horse serum for 10min. Subsequent incubations were carried out in the permeabilization buffer. Coverslips were incubated with an appropriate primary antibody and/or fluorescently conjugated phalloidin and/or fluorescently cpnjugated DNAse I for 60min at RT, washed 4 times in PBS and incubated with relevant fluorescently conjugated secondary antibodies at a concentration of 0.3µg/ml each for 60min at RT. Coverslips were then mounted in mounting medium (Southern), allowed to dry for 30min at RT and imaged on a Zeiss LSM710 microscope equipped with a standard set of lasers through a 63x oil apochromatic objective. Excitation wavelengths were 488, 543 and 633nm. Bandpass filters were set at 500–550 (AF488), 560–615 (AF568, AF594) and 650–720nm (AF647). Image acquisition was typically carried out at the 12-bit rate, in a “semi-blind” manner whereby the investigator could observe the signal in the AF488 or AF568/594 channel but not in the AF647 channel, thereby minimizing potential bias. Settings were optimized to ensure appropriate dynamic range, low background and sufficient signal/noise ratio.

### Quantification of synapse-specific levels of F-actin

Coverslips were processed as described above. To identify individual synapses, images were binarized in ImageJ using the “Moments” setting, and particles were counted automatically using the “Analyze Particles” command across the whole image excluding the cell body. Binarized data from the Homer channel was used for determination of synapses. To minimize contamination of the data with ROIs arising from non-specific staining or overlap of multiple synapses, only ROIs with areas ranging from 0.2 to 3μm^2^ were included in further analysis. All values of circularity were included in analysis. To measure the signal in the extrasynaptic regions in the vicinity of each synapse, ROIs defined by Homer puncta were subsequently expanded by 1pixel using the Enlarge functionin ImageJ. The values corresponding to the initial ROI, i.e. the synapse, were then subtracted from the respective values corresponding to the enlarged ROI, i.e. the synapse+its vicinity, yielding the values corresponding to the vicinity only. Background fluorescence intensity was measured in empty regions within the image for each channel and was subsequently subtracted from the signal.

### Statistics

Statistical analysis was carried out using GraphPad Prism5.0. Sample distribution was assessed using D’Agostino and Pearson’s omnibus normality test; to assess the significance of differences between datasets, two-tailed t test and 1-way ANOVA with Bonferroni’s multiple correction test were used in normally distributed datasets; otherwise, Mann-Whitney U test and 1-way ANOVA with Dunn’s post test were used. Box and whisker plots represent 10–90 percentile. Error bars in bar charts indicate standard error from the mean. ***P<0.001, **P<0.01, *<P<0.05.

---

**Supplementary Figure 1.**
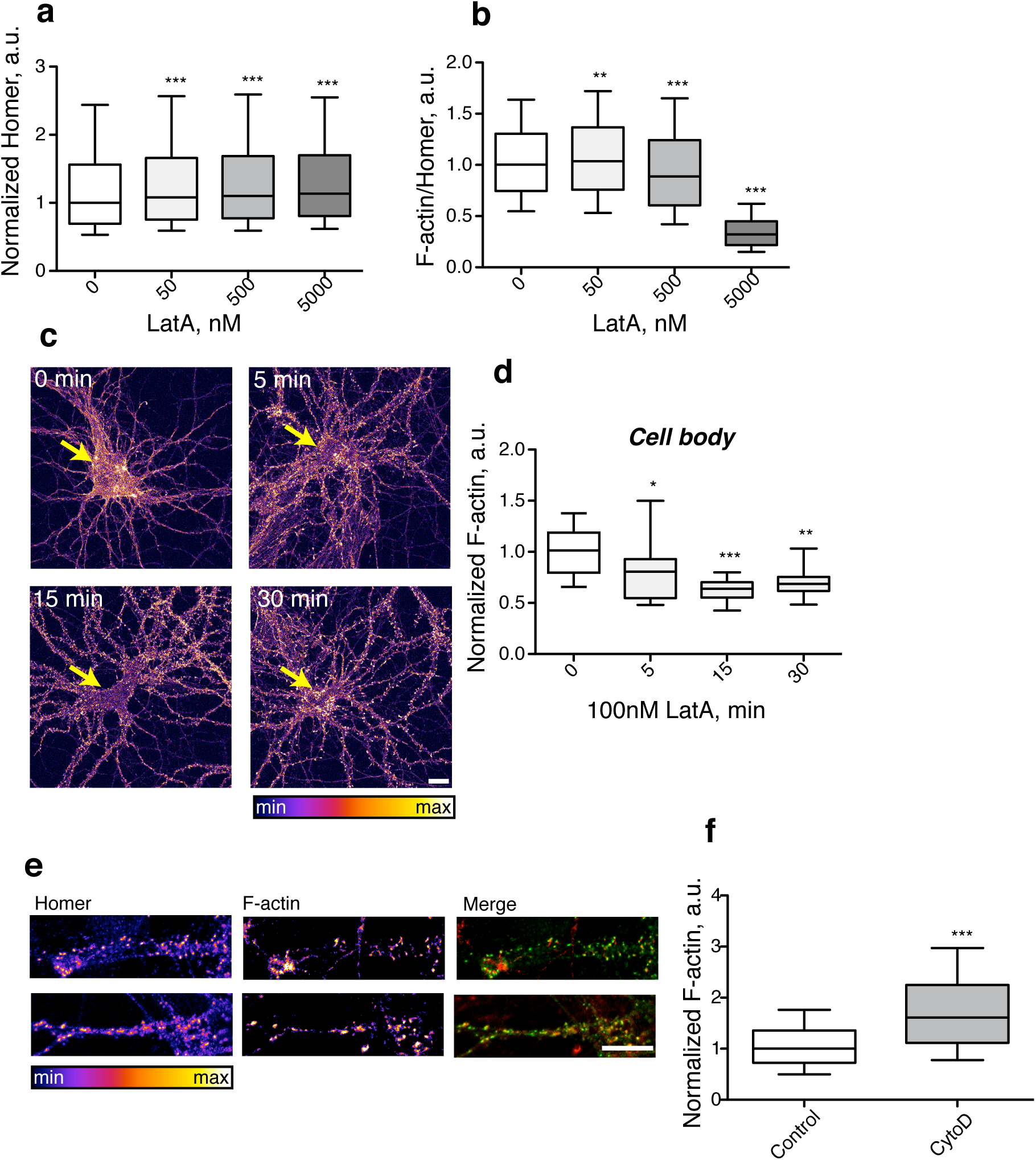
Additional data characterizing labile pool disruption. (**a**) Overnight inclubation with LatA results in an increase in the enlargement of the spine as evidenced by the increase in Homer intensity. N=3. (**b**) Enrichment of F-actin in the spine following overnight incubation with LatA. N=3. (**c**) Treatment with 100nM LatA results in a rapid loss of F-actin in the cell body of neurons. (**d**) Quantification of (c). N=5. (**e**) Treatment with 1uM CytoD results in a robust increase in spine actin. (**f**) Quantification of (e). N=4. Scale bar, 10um.

**Supplementary Figure 2.**
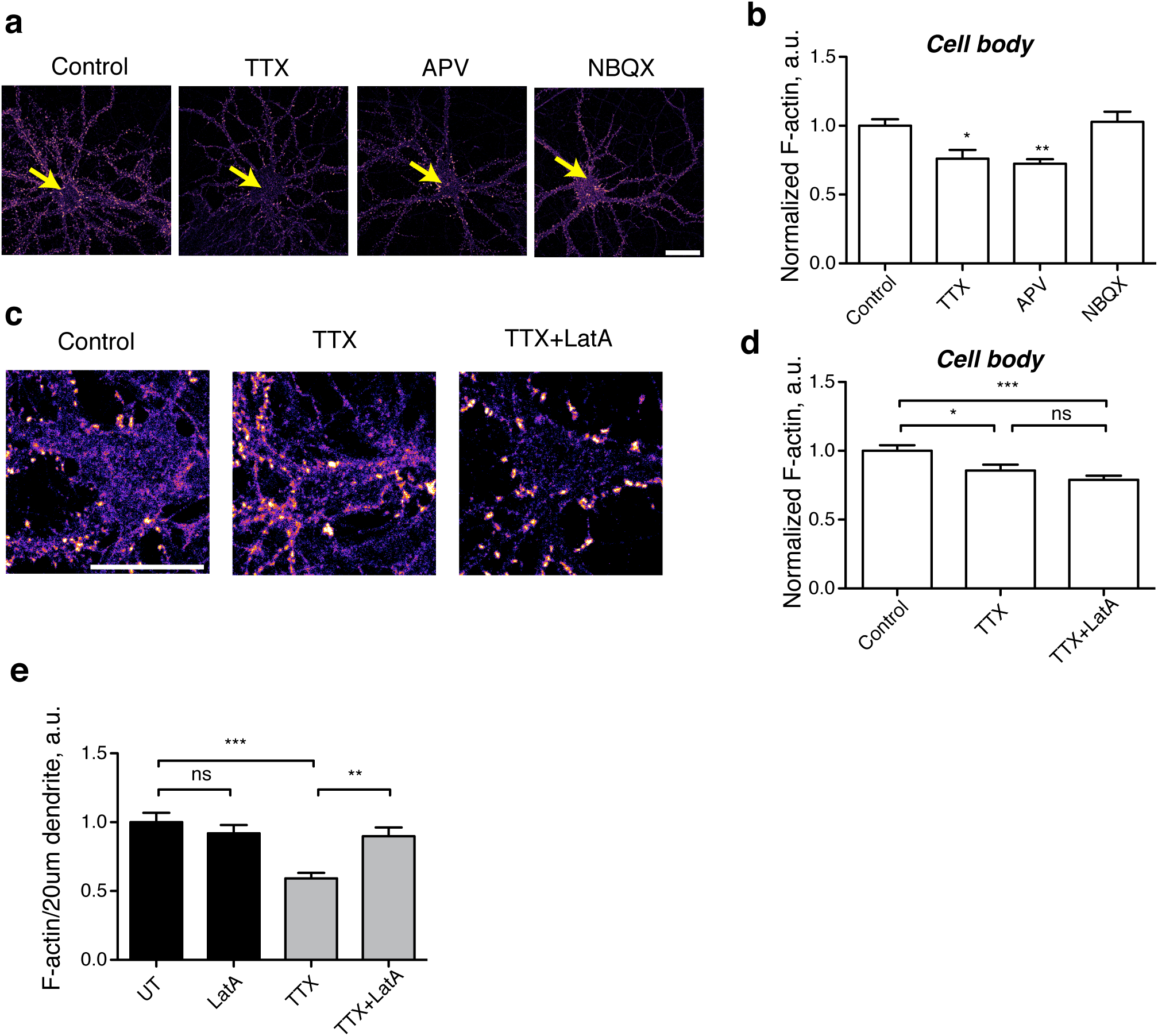
Additional data characterizing inactivity-induced depolymerization of spine F-actin. **(a)** Neurons were treated with indicated drugs for 48h, fixed and stained for F-actin. Arrows denote cell bodies. (**b**) Quantification of (a). N=3. (c) 30min treatment with LatA does not reverse the inactivity-induced loss of F-actin from the cell body. (d) Quantification of c. N=4. (**e**) 30min incubation with LatA significantly increases the amount of F-actin in the dendrites of TTX-treated, but not control neurons. N=4. Scale bar, 20um.

